# Hormone-dependent receptor docking controls calcium channel activity in plants

**DOI:** 10.64898/2026.07.10.737476

**Authors:** Vasyl Brykov, Lorena Huffer, Eva Medvecká, Tereza Korec-Podmanická, Daniela Kocourková, Lev Levenets, Karel Harant, Martina Schmidtová, Shiv Mani Dubey, Jana Krtková, Ivan Kulich, Roman Pleskot, Denisa Oulehlová, Matyáš Fendrych

## Abstract

The phytohormone auxin is a central coordinator of plant growth and development. Besides its canonical effect on gene transcription^1,2^, auxin triggers an ultra-rapid calcium ion influx that initiates the root gravitropic response^3^. The nature of the so-called rapid auxin pathway connecting the AFB1 auxin receptor^3,4^ and plasma membrane calcium channels remained unknown. Here, we show that auxin induces the direct interaction of the AFB1 receptor with the CNGC14 calcium channel. As the AFB1 receptor is independent of the ubiquitin ligase complex^5^, the auxin-induced interaction translates into relocalization of the receptor to the plasma membrane. We identify the interaction interface and provide evidence that the docking of the receptor to the channel complex activates Ca^2+^ influx and triggers growth inhibition. These findings position a calcium channel as an unprecedented component of the AFB1 auxin receptor complex. The ligand-dependent localization shift of a TIR1/AFB family receptor represents a novel paradigm in signal transduction and opens the possibility of unforeseen branches of auxin signaling pathways.

## Main

Plant roots navigate the heterogeneous soil to avoid obstacles, reach nutrient-rich spots, and anchor the plant. Root growth direction is steered by differential elongation of individual cells, e.g., during gravitropic bending. Gravitropic response is controlled by the transport and response to the phytohormone auxin, which inhibits root cell elongation^6^, so its redistribution leads to an elongation gradient and root bending. Inside the cell, auxin is perceived by the nuclear TIR1/AFB family receptors, where auxin functions as a molecular glue between the TIR1/AFBs and Aux/IAA co-receptor proteins that are the target-recognition components of the Skp1, Cullin, F-box-containing complex (SCF^TIR1/AFB^). Upon auxin binding, SCF^TIR1/AFB^ ubiquitinates the Aux/IAA proteins, which act as transcriptional repressors of auxin-responsive genes, and targets them for degradation, thereby changing the transcriptional landscape of the cell^1,2^.

However, in Arabidopsis, root gravitropic response is initiated by an ultra-rapid auxin-triggered growth inhibition that is steered by the AUXIN F-BOX 1 (AFB1) receptor^7–9^. Despite being closely related to the canonical TIR1, the AFB1 receptor functions independently from the SCF complex in the cytoplasm^3–5^, where it mediates ultrarapid changes of ion fluxes across the plasma membrane, calcium (Ca^2+^) influx being the hallmark of this response^3^.

The calcium channel responsible for this influx has been identified as the CYCLIC NUCLEOTIDE-GATED CHANNEL 14 (CNGC14), and the mutants lacking either the AFB1 or the CNGC14 lack the rapid auxin response^8,10^. Plant CNGCs are integral membrane proteins with the ion channel formed within a homo-or heterotetramer.

Individual CNGCs, out of the 20 CNGCs present in Arabidopsis, appear to be specific for particular processes and have different modes of activation^11,12^. The recent discovery of the cyclic nucleotide production by TIR1/AFB family receptors^13,14^ might provide a link to CNGC channel activation. However, plant CNGC channels turn out not to be gated by cyclic nucleotides^15^.

While the components of the pathway that triggers rapid Ca^2+^ influx in response to auxin have been identified, the molecular mechanism that connects the auxin receptor with a rapid change of plasma membrane ion fluxes remains enigmatic.

### Auxin receptor AFB1 associates with downstream effectors upon auxin perception

To reveal the mechanism of the AFB1 function, we generated a series of *Arabidopsis thaliana* lines expressing the orthogonal receptor ccvAFB1^16^. The modified (concave) ccvAFB1 binds exclusively the synthetic (convex) cvxIAA, while endogenous TIR1/AFB receptors cannot recognize the synthetic molecule^3^. Arabidopsis lines expressing a fluorescently tagged ccvAFB1 under the control of its native regulatory sequences triggered instantaneous Ca^2+^ influx and root growth inhibition in response to the treatment with cvxIAA in a vertical microfluidic imaging system (Fig. 1A-D, Movie S1). The orthogonal receptor-ligand pair thus fully reproduced the effects of the native indole-3-acetic acid auxin (IAA)^3^. Unlike the native auxin, the orthogonal ccvAFB1-cvxIAA system allowed us to bypass the complexity of the endogenous auxin signaling and to generate a sharp contrast between the active and inactive (cvxIAA-bound and cvxIAA-free form, respectively) AFB1 receptor *in planta*. Using transcriptomic profiling, we found that the action of ccvAFB1 was not associated with changes in gene expression that characterize the canonical SCF^TIR1/AFB^ activity, whereas an analogous ccvTIR1 did induce the expression of typical auxin-responsive genes when activated (Fig. S1A, Table S1). Interestingly, activated ccvAFB1 upregulated the expression of a single gene – TOUCH3, previously shown to be touch-and Ca^2+^-responsive^17,18^. This highlights that the AFB1 receptor acts by activating Ca^2+^ influx, not by inducing transcriptional changes.

**Fig. 1:**
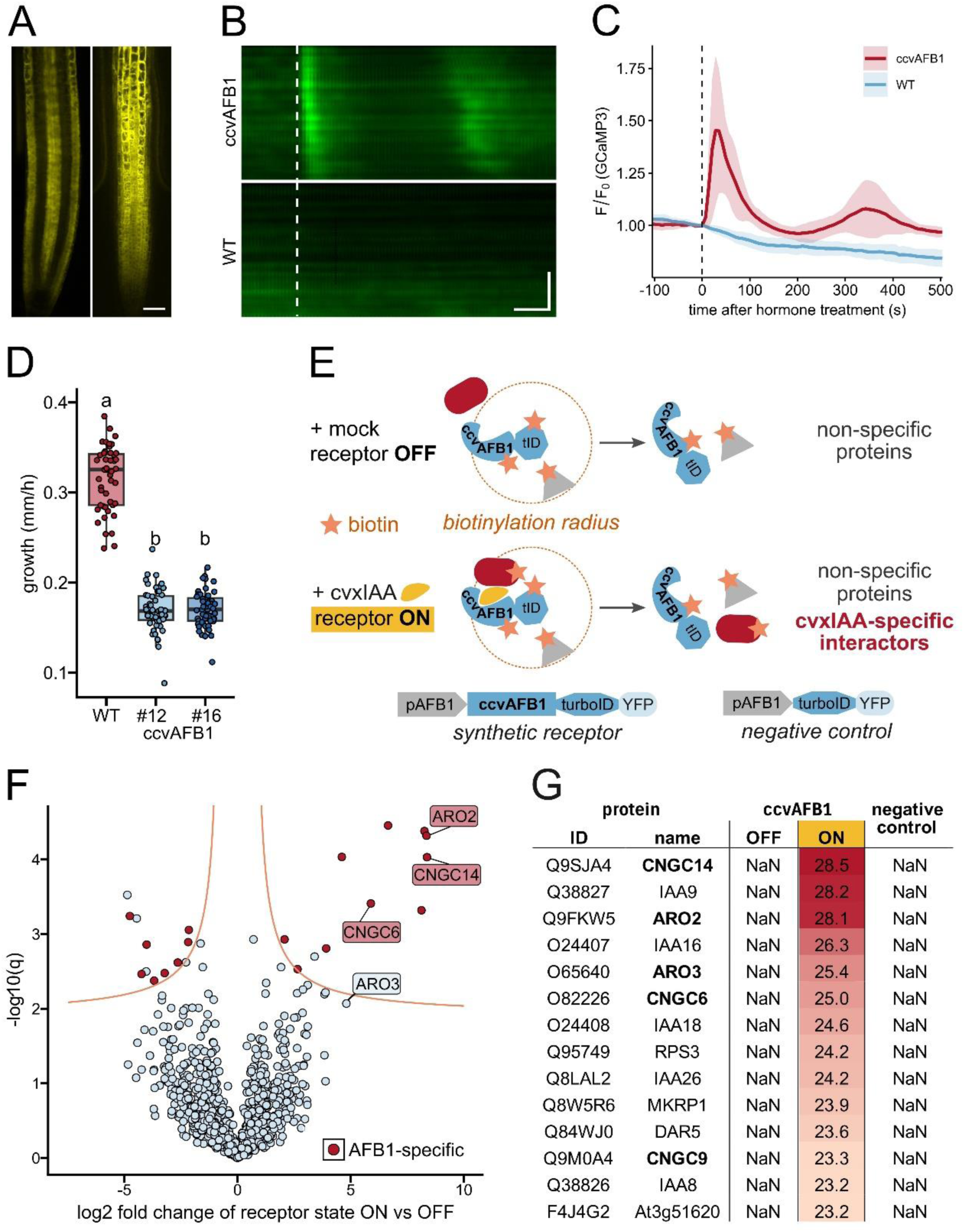
Active AFB1 associates with the ARO-CNGC complex. A) medial and epidermal optical sections of an Arabidopsis root tip expressing ccvAFB1-mVenus. Scale bar: 20 μm B, C) ccvAFB1-mScarlet triggers ligand-dependent cytosolic calcium ([Ca^2+^]cyt) transient. Representative kymograph (B) and quantification (C) of GCaMP3 intensity after application of 500 nM cvxIAA (dashed line) in the root elongation zone of control (WT) and ccvAFB1-expressing lines. In (C), GCaMP3 intensity was normalized per root to the pre-treatment time point. Scale bars: horizontal: 70s; vertical: 50 μm. D) cvxIAA inhibits root growth in ccvAFB1-expressing lines but not in the WT. A boxplot of root growth rates of WT (control) and two independent ccvAFB1-mScarlet lines treated with 500 nM cvxIAA, with all data points shown as dots. Statistical differences according to ordinary one-way ANOVA coupled with Tukey’s tests (p < 0.05) are indicated by letters. E) Schematic depiction of the proximity labeling experiment and the constructs used to generate Arabidopsis lines. F) Volcano plot showing the results of the ccvAFB1-turboID proxiome experiment. Orange lines represent the 0.2 false discovery rate threshold; proteins that pass the threshold and are absent in the negative control dataset are highlighted in red. G) List of specific interactors of active ccvAFB1. Proteins fulfill the following criteria: present in cvxIAA (ccvAFB1 ON), absent in mock (ccvAFB1 OFF), and absent in negative control, with a value over 23. Mean binary logarithms of protein intensity values are shown.

To elucidate how the AFB1 receptor triggers the downstream response, we searched for its interactors using co-immunoprecipitation (co-IP). However, this approach failed to reveal any interactors specific to the activated receptor. Interestingly, AFB1 co-immunoprecipitated and interacted with SKP1 proteins regardless of auxin presence (Fig. S1B, C, Table S2). We never detected CUL1, the ubiquitin ligase required for the SCF^TIR1/AFB^ activity^19^, and CUL1 localized solely to nuclei^20^ (Fig. S1D), excluding the interaction with AFB1 in the cytoplasm. This confirms that AFB1 operates fully independently from the SCF complex and the auxin transcriptional machinery^3,5^, while it can form a partial complex with SKP1A and SKP1B in the cytoplasm.

The co-IP results suggest that AFB1 interactions with its downstream effectors might be transient. Therefore, we opted for the proximity labeling approach^21,22^ and generated the ccvAFB1-turboID-YFP Arabidopsis line. We harnessed the binary nature of the ccvAFB1 orthogonal system and contrasted the set of proteins in the vicinity (proxiome) of the activated ccvAFB1 (cvxIAA-treated) against the proxiomes of the inactive form (mock-treated) as well as of the free TurboID-YFP control (Fig. 1E) from Arabidopsis roots. This approach yielded a short list of fully specific, ligand-dependent ccvAFB1 interactors (Fig. 1F, G, Table S3). Remarkably, the active AFB1 proxiome contained CNGC channels, ARMADILLO-REPEAT ONLY proteins (AROs), and we also revealed several Aux/IAAs proteins (Fig. 1F). The most abundant interactor of the active AFB1 was CNCG14, a PM-localized Ca^2+^ channel required for the auxin-induced Ca^2+^ transient^10^. Plant CNGCs form heterotetramers at the plasma membrane^15^, and CNGC6, 9, and 14 appearing in our dataset are expressed in Arabidopsis roots^23^. ARO proteins have recently been identified as components of the CNGC channel complex and are required for CNGC activity^24^. The proxiome of active AFB1 strongly suggests that upon binding the ligand, AFB1 directly associates with the CNGC-ARO channel complex to activate the auxin-induced Ca^2+^ influx and the downstream root growth inhibition response.

### Auxin receptor is recruited to the plasma membrane in a ligand-dependent manner

How can a cytoplasmic receptor encounter a plasma membrane (PM) localized ion channel complex? We re-visited the subcellular localization of the native AFB1 receptor in root epidermal cells using high-resolution spinning disk microscopy.

Above the background of the cytoplasmic signal, we detected AFB1 signal foci that localized to the cellular periphery (Fig. 2A, B). The treatment with nanomolar concentrations of the natural auxin (IAA) significantly increased the recruitment of AFB1 into the foci (Fig. 2A-C).

**Fig. 2:**
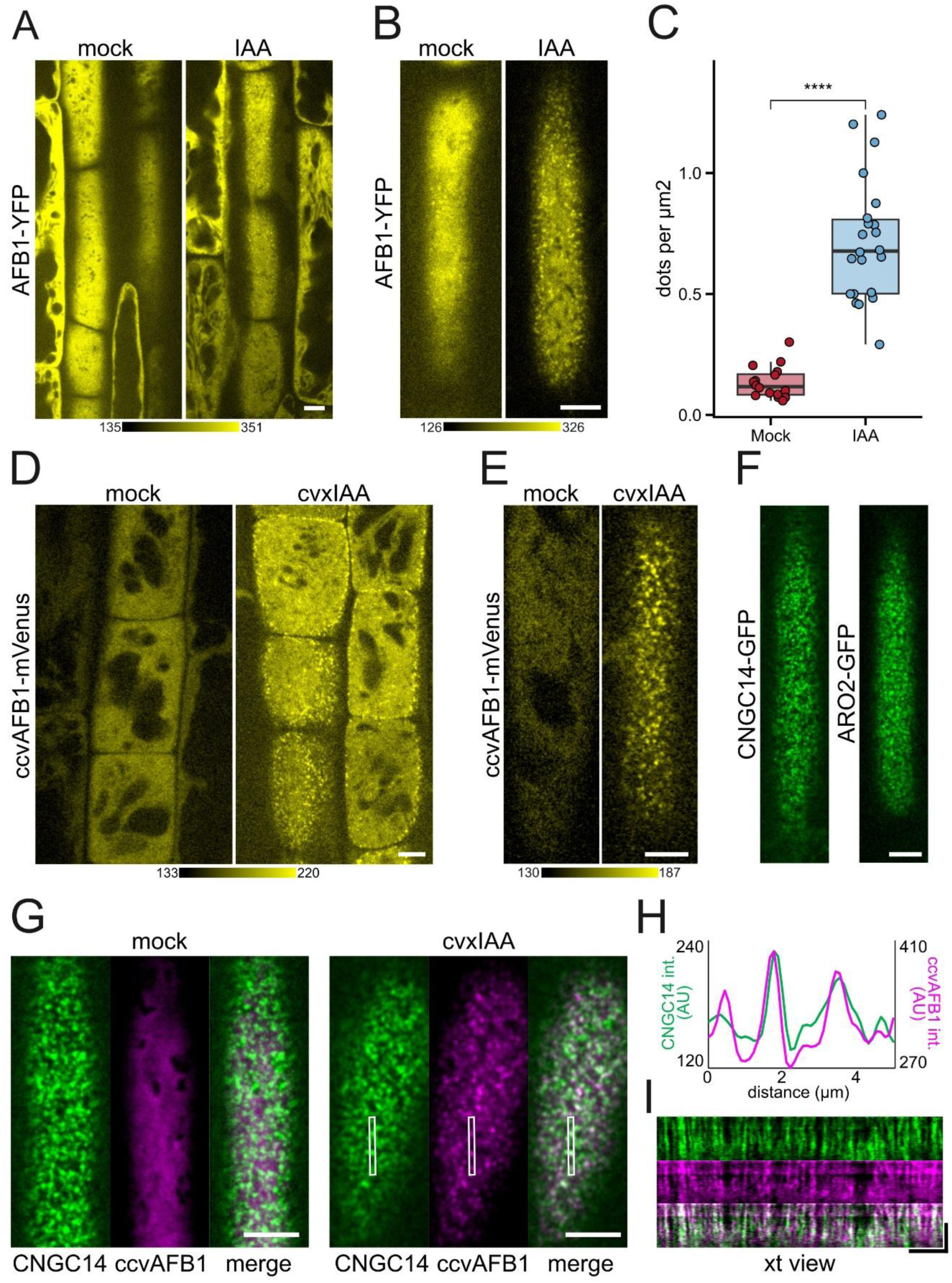
AFB1 receptor docks at the plasma membrane upon hormone perception. A-C) Auxin (100 nM IAA) stimulates recruitment of AFB1-YFP into PM-localized foci in root epidermal cells. A) overview image of root transition zone, B) tangential image of a single cell surface, C) quantification of PM foci density. **** indicates P<0.0001 according to ordinary one-way ANOVA. D, E) ccvAFB1-mVenus shows a binary localization switch upon perception of the cvxIAA ligand. D) optical section of root transition zone shows ccvAFB1 foci at cell periphery. E) tangential imaging of a single cell surface. Treatments with mock or 500 nM cvxIAA are indicated. F) CNGC14-GFP and ARO2-GFP localize to PM foci. Tangential images of root epidermal cell surface. G-I) cvxIAA (500 nM) induced the colocalization of ccvAFB1-mScarlet with CNGC14-GFP in root epidermal cells. G) Single cell surfaces of Arabidopsis roots expressing CNGC14-GFP (green) and ccvAFB1-mScarlet (magenta). H) A plot of CNGC14-GFP and ccvAFB1 fluorescence intensities along the line highlighted in (G). I) The kymograph (time-space view, xt view) shows the colocalization of ccvAFB1-mScarlet (magenta) with CNGC14-GFP (green) in PM foci upon cvxIAA treatment. Horizontal dimension represents the x-dimension, while horizontal dimension represents t-time. Scale bars in all figures = 5 μm; vertical scale bar in(I) = 60 s.

To test whether the hormone ligand is necessary to trigger the PM recruitment, we analyzed the localization of the synthetic ccvAFB1 in response to cvxIAA. Whereas the signal of inactive ccvAFB1 was purely cytoplasmic, addition of the ligand caused a clear recruitment into foci (Fig. 2D, E; Movie S2). Both cross-section and tangential imaging of the cellular surface revealed the localization of the foci to the PM. This binary response confirms that ccvAFB1 approaches the plasma membrane only upon perception of the cvxIAA ligand, which does not occur in the natural system.

Meanwhile, the native AFB1 receptor localizes to PM foci in response to the endogenous auxin, and additional auxin increases its PM recruitment.

The localization of AFB1 to PM foci prompted us to compare it to the localization of its top interactors, CNGC14 and ARO2. CNGC14 and ARO2 formed PM foci that highly resembled the AFB1 foci (Fig. 2F). We crossed ccvAFB1-mScarlet with either native-promoter driven GFP-tagged CNGC14 or ARO2 to test whether AFB1 is being recruited to the same PM foci. Without the ligand, in root epidermal cells, CNGC14 localized to PM foci, and ccvAFB1 showed a cytoplasmic signal (Fig. 2G). Upon treatment with cvxIAA, ccvAFB1 largely colocalized with CNGC14, as determined using intensity plot profiles and kymograph analysis (Fig. 2G-I, Movie S3). Similarly, ccvAFB1 colocalized with ARO2 only upon cvxIAA treatment (Fig. S2A, Movie S4).

These results show that the AFB1 receptor undergoes a binary localization switch between a purely cytoplasmic one in the inactive state and docked to the CNGC14-ARO complex upon perception of its hormone ligand.

### Receptor docking at the plasma membrane depends on the CNGC14 channel

The TIR1/AFB proteins were shown to possess adenylate cyclase (AC) and guanylate cyclase (GC) activity that is stimulated by the formation of TIR1/AFB-auxin-Aux/IAA complex^13,25^, and it was suggested that the cyclic GMP (cGMP) might activate CNGC channels^25^. We analyzed ligand-dependent cellular Ca^2+^ transients in the lines expressing either control ccvAFB1 or ccvAFB1dAC/GC receptors carrying mutations in both putative AC and GC domains which were previously shown to impede the AC and GC activity, respectively^13,25^. In both the control and AC/GC-deficient lines, the cvxIAA ligand triggered an instantaneous Ca^2+^ transient (Fig. 3A, Fig. S3A-C, Movie S5), and the ccvAFB1dAC/GC receptor was still recruited into PM foci (Fig. 3B), indicating that these mutations do not interfere with the receptor activity and activation of CNGC14-mediated Ca^2+^ influx. This result aligns with the finding that plant CNGCs are generally not regulated by cyclic nucleotides^15^.

**Fig. 3:**
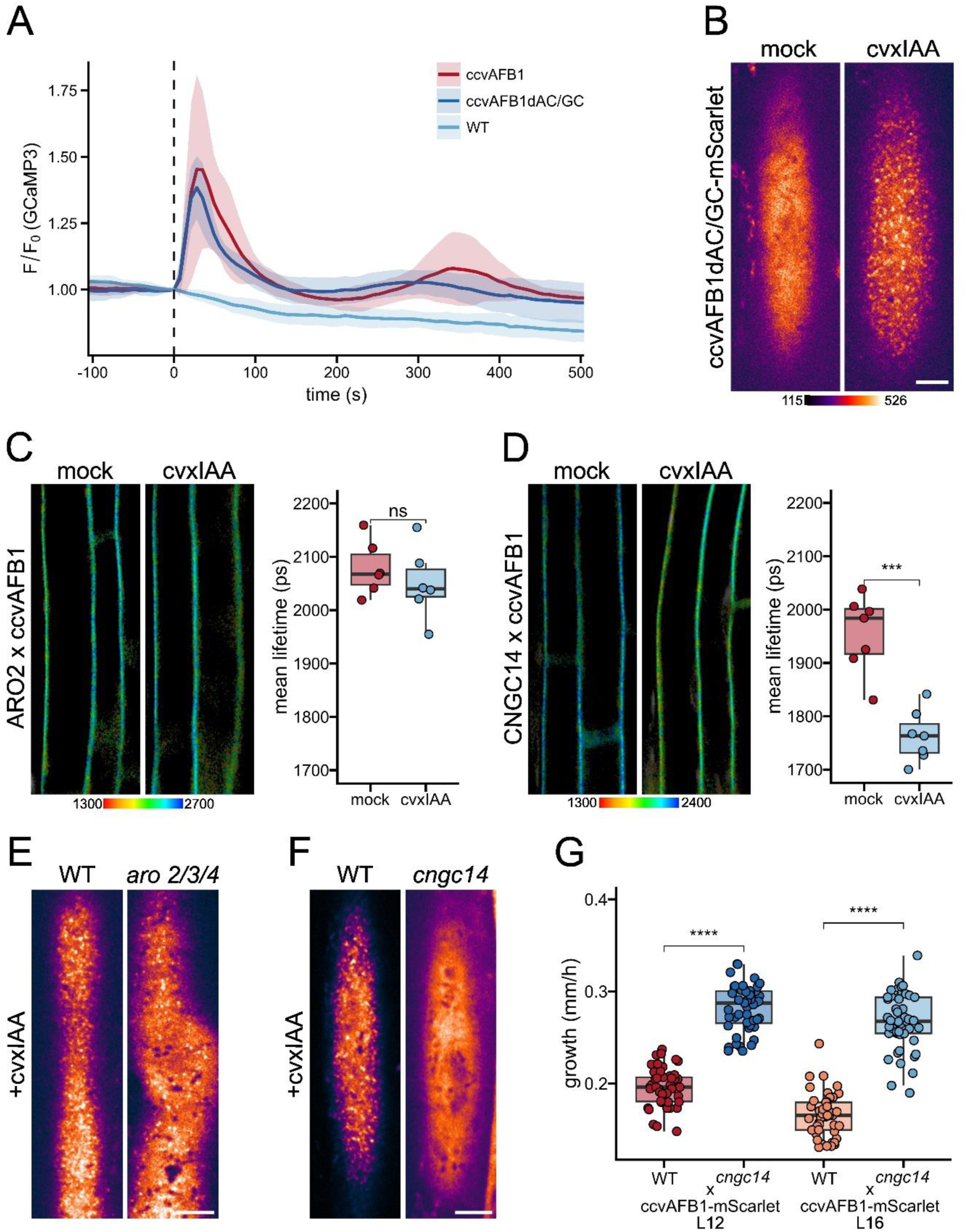
CNGC14 is required for docking of the receptor at the plasma membrane. A) ccvAFB1dAC/GC receptor triggers an instantaneous cytosolic calcium transient in response to the cvxIAA ligand. Quantification of the normalized GCaMP3 fluorescence intensity (F/F0 GCaMP3) in root epidermal cells of ccvAFB1, ccvAFB1dAC/GC, and WT lines in response to 500 nM ccvIAA (time point 0, dashed line). GCaMP3 intensity normalized per root to the pre-treatment timepoint intensity. Shaded areas correspond to standard deviation. B) ccvAFB1dAC/GC-mScarlet receptor relocalizes to PM foci upon cvxIAA (500 nM) treatment. Scale bar: 5 μm. C) *In vivo* fluorescence lifetime imaging and quantification of ARO2-GFP x ccvAFB1-mScarlet in root epidermal cells. D) *In vivo* fluorescence lifetime imaging and quantification of CNGC14-GFP x ccvAFB1-mScarlet in root epidermal cells shows that the lifetime of CNGC14-GFP is reduced significantly in the presence of ccvAFB1-mScarlet after addition of the cvxIAA ligand. In C, D), the pseudocolor represents the mean fluorescence lifetime, the intensity represents the photon count. The pseudocolor key shows the scale of the fluorescent lifetimes. A representative lifetime image and quantifications are shown. The quantification shows the mean lifetimes of several roots averaged and compared between mock and cvxIAA treatment. Statistical differences according to one-way ANOVA are indicated by stars. E) ccvAFB1 forms PM foci in the *aro2/3/4* mutant background. Details of the cell surface of WT and *aro2/3/4* lines expressing the receptor treated with 500 nM cvxIAA. Scale bar: 5 μm. F) ccvAFB1 is unable to form PM foci in the *cngc14* mutant. Details of the cell surface of WT and *cngc14* lines expressing the receptor treated with 500 nM cvxIAA. Scale bar: 5 μm. G) In the *cngc14* mutant, the ccvAFB1-mScarlet receptor fails to inhibit root growth in response to cvxIAA. Box plot of root growth rates of two independent ccvAFB1-mScarlet lines crossed into *cngc14* mutant treated with 500 nM cvxIAA; outcrossed mutant and control lines are compared. Statistical differences according to one-way ANOVA are indicated by stars.

We therefore hypothesized that a direct auxin-dependent interaction of AFB1 with either CNGC14 or AROs recruits the receptor to the PM, which leads to CNGC14 channel activation. To test this, we employed Förster Resonance Energy Transfer - Fluorescence Lifetime Imaging Microscopy (FRET-FLIM) between native-promoter driven CNGC14-GFP or ARO2-GFP donor and ccvAFB1-mScarlet acceptor in Arabidopsis roots. Addition of the cvxIAA ligand did not change the ARO2 or CNGC14 lifetime in the absence of ccvAFB1 (Fig. S3D, E). The ARO2-GFP fluorescence lifetime was not significantly influenced by ccvAFB1-mScarlet regardless of the hormone ligand presence (Fig. 3C), although a trend towards lower lifetimes was visible after cvxIAA treatment. On the other hand, the CNGC14-GFP showed a significant, ligand-dependent lifetime decrease in the presence of the ccvAFB1 receptor (Fig. 3D), indicating that AFB1 interacts with CNGC14 upon binding the hormone, and confirming the ligand-dependent colocalization data with an independent fluorescence spectroscopy method^26^. In addition, the ligand-dependent AFB1-CNGC14 interaction revealed by FLIM imaging confirms and functionally specifies the proxiome results.

Next, we analyzed ccvAFB1 PM recruitment in the *aro2/3/4* and *cngc14* mutants, which both lack the rapid auxin-induced Ca^2+^ influx and growth inhibition, similarly to the *afb1* mutant^3,10,24^. While the ccvAFB1-mScarlet PM foci still formed in a ligand-dependent manner in the *aro2/3/4* mutant (Fig. 3E), the PM recruitment of the AFB1 receptor was fully abolished in the *cngc14* mutant (Fig. 3F). If AFB1 docking to CNGC14 is needed for the activation of the auxin response, the ccvAFB1 effect should be lost in the absence of CNGC14. We therefore tested the cvxIAA-induced root growth inhibition - the direct *in vivo* readout of the AFB1 pathway - in ccvAFB1 crossed to the *cngc14* mutant background. While cvxIAA application inhibited root growth in the outcrossed wild type plants, the ccvAFB1 receptor failed to inhibit root growth in the *cngc14* mutant background (Fig. 3G). These results demonstrate that ARO proteins are not needed for the PM docking of active AFB1, even though AROs are required for the CNGC channel functionality^24^. Instead, the AFB1 receptor is recruited to the CNGC14-ARO channel complex in a strictly CNGC14-dependent manner, and this interaction triggers the CNGC14 activity that results in root growth inhibition.

### Auxin bridges the AFB1 receptor with the C-terminal helix of CNGC14

To identify the mechanism of the ligand-dependent AFB1-CNGC14 interaction, we employed structural modelling using a local installation of the Alphafold3 (AF3) program^27^. We explored two possible scenarios: a) AFB1 - auxin (IAA) - IAA9 - CNGC14 tetramer, and b) AFB1 - auxin (IAA) - CNGC14 tetramer. We included IAA9 as the most abundant Aux/IAA protein detected in the active AFB1 proxiome (Fig. 1G). The scenario including IAA9 yielded a highly confident interaction between IAA9-IAA-AFB1, which corresponds to the experimentally resolved interaction between the Aux/IAA polypeptide, IAA, and TIR1^28^ (Fig. S4A). Nevertheless, even using several different seeds for structural modelling did not result in a confident position of IAA9 or AFB1 towards the CNGC14 tetramer. Omitting IAA9 from the modelling, however, revealed an interaction site between a C-terminal helix of CNGC14 (amino acid residues 683-707) and the auxin binding pocket of AFB1, with a very low predicted alignment error (PAE), corresponding to the reliability of experimental methods^27^ (Fig. 4A-B). Importantly, the interaction was predicted only in the presence of the IAA ligand (Fig. 4B). Consistent with our FRET-FLIM data, the AF3-based structural model of the IAA-mediated AFB1-CNGC14 interaction positions the C-terminus of both proteins in proximity, thus allowing FRET (Fig. 4A). Similar to the IAA-mediated TIR1-Aux/IAA interaction, our model suggests that the auxin molecule functions as a molecular glue bridging the AFB1 and the C-terminal part of CNGC14 (Fig. 4C).

**Fig. 4:**
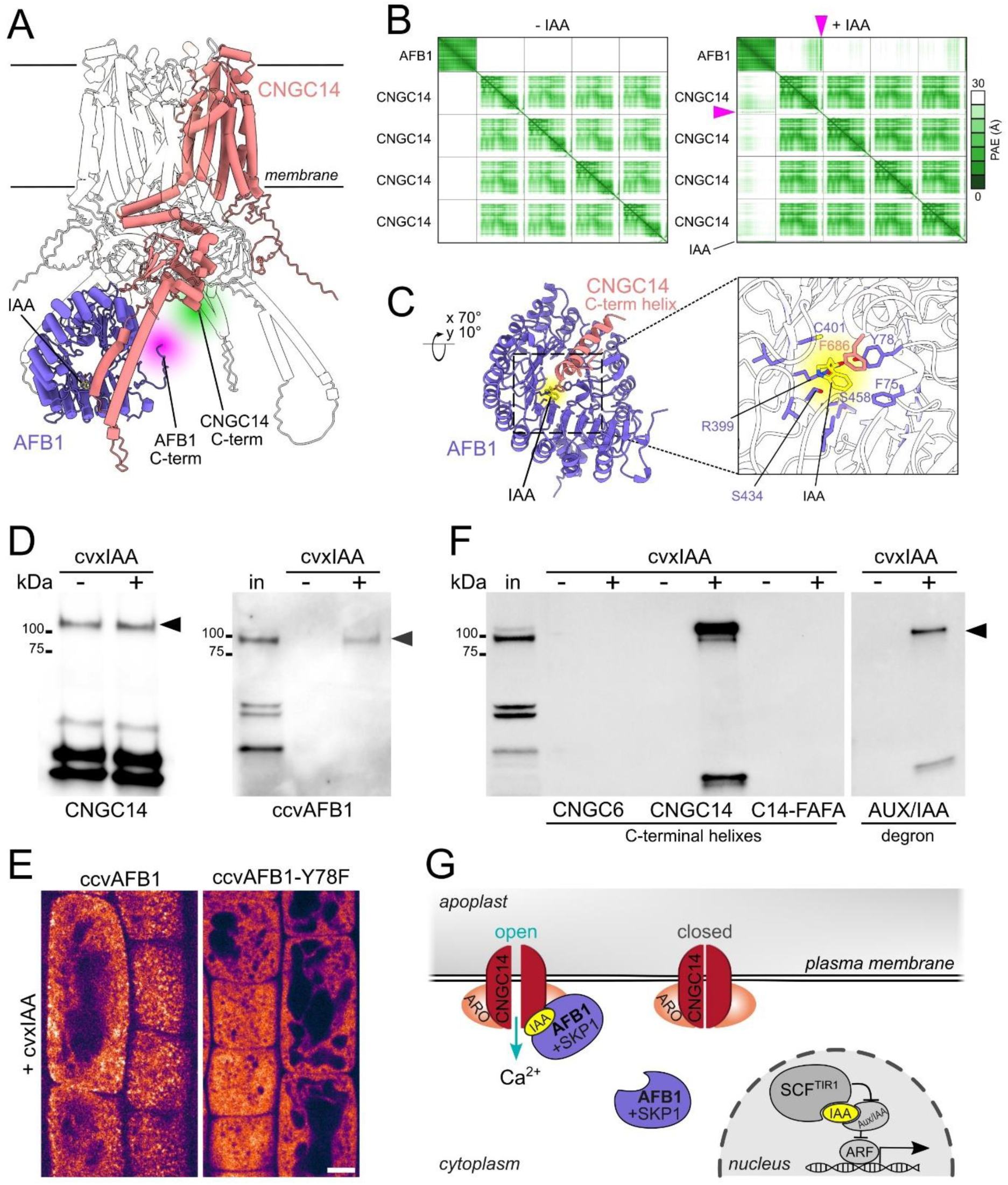
AFB1 interacts with the CNGC14 through its auxin-binding pocket. A) AlphaFold3 prediction of the IAA-dependent interaction between AFB1 and the CNGC14 tetramer. AFB1 is represented in blue using a ribbon structure, while the AFB1-interacting chain of the CNGC14 tetramer is shown in light red. The other three chains are displayed in a transparent ribbon representation. IAA is depicted in yellow using a licorice representation. The positions of the C-termini of AFB1 and CNGC14 are indicated in green and purple, respectively. B) The predicted alignment error (PAE) plots for the complex composed of AFB1 and the CNGC14 tetramer with and without IAA. The plot shows a confidently predicted interaction between AFB1 and CNGC14 mediated by the IAA molecule. C) AlphaFold3 prediction of the IAA-dependent interaction between AFB1 (blue) and the C-terminal helix of CNGC14 (amino acids 683-708, light red). The IAA molecule is shown in yellow using a licorice representation. The zoomed-in picture on the right illustrates a binding pocket formed by AFB1 and the C-terminal helix of CNGC14. Amino acid residues that coordinate IAA are shown in a licorice representation. D) ccvAFB1 co-immunoprecipitates with CNGC14 exclusively in the presence of cvxIAA. CNGC14-GFP purified from tobacco leaves detected by anti-GFP (left) traps ccvAFB1-mScarlet detected by anti-mCherry from the Arabidopsis total protein extract after the addition of cvxIAA (right). Minus cvxIAA = mock (DMSO), plus cvxIAA = treatment with 50 μM cvxIAA, in = input. E) The 78Y residue within the auxin binding pocket of AFB1 is crucial for the PM recruitment. Left: ccvAFB1-mScarlet driven by a weak TIR1 promoter decorates PM foci in root cells upon 500 nM cvxIAA. Right: pTIR1:ccvAFB1-mScarlet with a Y78F substitution remains cytoplasmic. Scale bar: 5 μm. F) ccvAFB1 from the Arabidopsis total protein extract co-immunoprecipitates with the CNGC14 C-terminal helix in an cvxIAA-dependent manner; the mutated CNGC14-C-terminal FAFA helix and the C-terminal helix of CNGC6 show no interaction either with or without cvxIAA. ccvAFB1-mScarlet was detected by anti-mCherry. On the right, the cvxIAA-dependent ccvAFB1 recruitment by the canonical Aux/IAA peptide is shown. Minus cvxIAA = mock (DMSO), plus cvxIAA = treatment with 50 μM cvxIAA during the pull-down assay, in = input of the total protein extract. G) A conceptual model of the rapid auxin signaling pathway. At low auxin levels, the AFB1 receptor (blue) remains cytoplasmic. When the auxin (IAA in yellow) concentration increases, AFB1 docks to the CNGC14-ARO complex (CNGC channel in red, ARO in orange) and facilitates Ca^2+^ influx, subsequently leading to root growth inhibition. Meanwhile in the nucleus, auxin is perceived by the canonical TIR1/AFB receptors that form the ubiquitin-ligase complex (SCF^TIR1^) to modulate transcription of auxin responsive genes.

To confirm the predicted direct AFB1-CNGC14 ligand-dependent interaction experimentally *in vitro*, we expressed CNGC14-GFP heterologously in the leaves of *Nicotiana benthamiana*. We used the purified CNGC14-GFP protein to pull down ccvAFB1-mScarlet from the Arabidopsis total protein extract. In accordance with the previous results, ccvAFB1 was co-immunoprecipitated with CNGC14-GFP upon addition of the cvxIAA ligand, while it was absent in the control condition (Fig. 4D). This result demonstrates that the CNGC14-AFB1 complex assembles specifically in the presence of auxin, and its formation is robust enough to occur *in vitro* in the context of a total protein extract.

To test whether other proteins participate in the assembly of the complex; we analyzed proteins co-immunoprecipitated with CNGC14 (Fig. 4D) using mass spectrometry. We identified three proteins enriched specifically upon cvxIAA treatment: AFB1, SKP1A, and SKP1B (Fig. S4C, Table S6), confirming that AFB1 forms a stable complex with SKP1 (Fig. S1B, C). SKP1 cannot directly mediate the auxin-dependent CNGC14-AFB1 interaction, as it interacts with the N-terminal F-box domain of TIR1/AFB proteins^28^, which is distant from the auxin binding pocket of AFB1. Importantly, Aux/IAA proteins were not detected, confirming our structural modeling results. Aux/IAAs are known nuclear proteins^29,30^, their presence in the AFB1 proxiome thus probably reflects their interaction with AFB1 in the nucleus^29,30^, while they don’t participate in the CNGC14-AFB1 complex at the PM.

The modelling pointed to the auxin binding pocket as the interface between the receptor and the channel. In Arabidopsis, AFB1 is specialized for triggering the ultra-rapid Ca^2+^ transient^3^. We examined the amino acid conservation around the auxin-binding pocket and found that Arabidopsis AFB1 carries an F78Y mutation (Fig. S4B), which lies at the CNGC14-AFB1 contact interface (Fig. 4C). We expressed the ccvAFB1-Y78F under the control of a weak promoter to obtain a good signal-to-noise image of the PM foci. The ccvAFB1-Y78F retained the ability to bind auxin and form the Aux/IAA coreceptor complex (Fig. S4D)^31^. However, unlike the control protein that showed the ligand-dependent PM foci recruitment, ccvAFB1-Y78F remained cytoplasmic (Fig. 4E). Accordingly, the ccvAFB1-Y78F receptor failed to inhibit root growth even when expressed from a strong promoter (Fig. S4E), while the control lines inhibited root growth in a ligand-dependent manner.

Further, to experimentally validate the CNGC14 interaction interface, we immobilized the CNGC14 C-terminal helix (amino acid residues 679-707) to agarose beads and incubated it with total protein extract from ccvAFB1-mScarlet-expressing Arabidopsis plants. As a negative control, we prepared an analogous domain from CNGC6, which was identified in the active AFB1 proxiome (amino acid residues 712-740). The ccvAFB1 showed a strong cvxIAA-dependent recruitment to the CNGC14 C-terminal helix *in vitro* (Fig. 4F), while we observed no binding to CNGC6. Finally, we mutated two phenylalanine residues (F686>A and F695>A) in the CNGC14 C-terminal helix that we identified as crucial for the interaction, as they face auxin and the AFB1 auxin binding pocket (Fig. 4C, S4F). The resulting CNGC14-C-terminal FAFA helix fully lost the ability to recruit ccvAFB1 in response to the cvxIAA ligand (Fig. 4F).

The cvxIAA-driven formation of CNGC14-AFB1 complex was as strong and specific as the cvxIAA-driven formation of the canonical Aux/IAA-AFB1 coreceptor^1,2,31^ (Fig. 4F). In summary, these results identify the AFB1 auxin-binding pocket and the CNGC14 C-terminal helix as the auxin-dependent interaction interface of the auxin receptor and the calcium channel.

## Conclusions

Our results show that auxin mediates a direct and specific interaction between the AFB1 receptor and the C-terminal part of the CNGC14 channel (Fig. 4G), analogously to the formation of the TIR1/AFB-auxin-Aux/IAA co-receptor complex^1,2^. This finding positions the CNGC14 channel as a novel AFB1 auxin co-receptor.

Further, this opens the intriguing possibility that auxin can act as a molecular glue between TIR1/AFB family F-box receptors and so far undiscovered substrates, opening potential unforeseen branches of the auxin signaling pathways. In the nuclear auxin pathway, TIR1/AFB substrate recognition triggers proteasomal degradation of the target co-receptor^1,2,28,32^. AFB1 is unable to assemble into the ubiquitin ligase complex^5^, and therefore the substrate recognition does not lead to ubiquitinylation, but instead translates into the localization shift of the AFB1 receptor through docking to the CNGC14 complex, leading to activation of Ca^2+^ influx.

The mechanism of plant CNGC channel gating remains unknown. The activity of CNGC channels is generally regulated by interaction with ARO proteins^24^ and specifically by phosphorylation^11^, by interactions of their C-terminal domains with calmodulins, and by other modes^33^. Calmodulins bind to the IQ peptide and were shown to both activate and inhibit CNGC channels^34–36^. Given that the AFB1 binding site is adjacent to the IQ peptide, AFB1 docking may change the mutual position of the C-terminal domains of CNGC14 channel monomers, leading to channel opening. Alternatively, or in parallel, docking of AFB1 to CNGC14 might displace calmodulin proteins, due to the close proximity of the IQ peptide and the AFB1-binding region^34,37,38^. The fact that we revealed no calmodulin proteins in the active AFB1 proxiome supports this hypothesis; on the other hand, we did not observe differential calmodulin abundance on the purified CNGC14 channel upon ccvAFB1 binding (Fig. S4C). The exact mechanistic insight of how AFB1-CNGC14 interaction activates the channel remains to be determined. Nevertheless, we provide compelling evidence that this interaction triggers Ca^2+^ influx into cells, which triggers the downstream root growth inhibition^39^.

In summary, the subcellular relocalization of a hormone receptor and its direct association with the downstream effector - a calcium channel - represents an unprecedented paradigm of hormone signal transduction in eukaryotic cells.

## Methods

### Molecular cloning

Cloning procedures were carried out using the GoldenBraid system^1^. All the generated constructs and transgenic lines are described in Table S4.

The turboID-YFP tag was amplified with either 762+765 primers for domestication of the free turboID control or 764+765 primers for domestication of the tag for C-terminal fusion, see Table S5 for primers used. Addgene vector was used as a template.

The SKP1A coding sequence was amplified using primers 879+880, CUL1 using primers 881+882 and 919+920 using genomic *Arabidopsis thaliana* DNA as a template (Table S5). The pAFB1 promoter (958 bp genomic fragment upstream of the AT4G03190 start codon) was amplified using primers 772 and 773; the AFB1 terminator (411 bp genomic fragment downstream of the AT4G03190 stop codon) was amplified using primers 992 and 993 (Table S5). The ccvAFB1dACGC mutant version was created by changing E549 (AC motif), and G517 (GC motif) to alanine using the primers 970-973 (Table S5), the ccvAFB1^2^ was used as a template.

The ccvAFB1-Y78F mutant variant was generated by amplifying the ccvAFB1 sequence using the primers 187 and 189, and then combining it with the rest of the ccvAFB1 sequence^2^.

### Plant material and growth conditions

A complete list of the newly generated and previously established lines is provided in Table S4. For stable plant transformation we followed a floral dip protocol described earlier^3^, and the *Arabidopsis thaliana* (L.) Heynh. ecotype Columbia-0 (Col-0) was used as the background, unless stated otherwise. To achieve the transient expression in *Nicotiana benthamiana* leaves, we followed a protocol described earlier^4^.

To analyze the colocalization and the FRET-FLIM imaging, we crossed the pARO2:ARO2-GFP, the pARO3:ARO3-GFP^5^, and the pCNGC14:CNGC14-GFP^6^ lines with the pAFB1:ccvAFB1-mScarlet line. Experiments were carried out in the F1 generation. In Fig. 3F, the ccvAFB1-mScarlet lines were crossed with *cngc14-1*, the wild type and homozygous background were selected in the F2 generation by genotyping^6^.

Seeds for all experiments were surface sterilized with chlorine gas, and sown on growth medium (½ Murashige and Skoog basal salt mixture supplemented with 1% sucrose and 0.5gL-1 MES, pH 5.7, 1% plant agar, Duchefa Biochemie). Seeds were stratified for 2 days at 4 °C, then the seedlings were cultivated vertically for 4-5 days at 23°C and 16/8 day/night with ∼100 μmol m-2 s-1 photon flux density.

### Root growth assay

For the root growth rate test, seedlings were transferred to plates filled with growth medium containing treatment or mock of indicated concentration and scanned using the Perfection V700 flatbed scanner (Epson) immediately and then after the indicated treatment time. Root length increment was determined using the Segmented line tool in the FIJI software^7^. The root growth response to cvxIAA was calculated as the length increment in treatment divided by the length increment in control conditions.

### Pull-down experiments

#### AFB1-mCitrine interactors

Protein pull-down assays were conducted to identify AFB1-interacting proteins from *Arabidopsis thaliana* whole seedlings and roots. Seeds of pAFB1:AFB1-mCitrine^8^ were grown for 6 days and then treated on plates with either 100 nM indole-3-acetic acid (IAA) or ethanol (mock control) in water for 2 minutes. Approximately 200 mg of seedlings or roots were flash-frozen in liquid nitrogen and ground to a fine powder. Proteins were extracted in a buffer containing 100 mM Tris-HCl (pH 7.5), 150 mM NaCl, 0.5% NP-40, 1 mM PMSF, and cOmplete protease inhibitor cocktail (Roche). Extraction was performed at 4 °C for 40 minutes, followed by centrifugation at 14,000 rpm for 20 minutes at 4 °C to remove debris. The cleared protein extracts were incubated with GFP-Trap agarose beads (Chromotek), which were pre-equilibrated in the extraction buffer. Both equilibration and subsequent washing steps involved centrifugation at 450 × g for 1 minute to pellet the beads. Binding was performed for 1 hour at 4 °C under gentle rotation. Beads were washed twice with the wash buffer (100 mM Tris-HCl, pH 7.5; 150 mM NaCl; 0.5% NP-40) and twice in the wash buffer without detergent. After the final wash, beads were stored at −80 °C until further analysis.

#### CNGC14-GFP - AFB1mScarlet pulldown

As the source of CNGC14, we used leaves of *Nicotiana benthamiana* transiently expressing 35S:CNGC14-GFP together with UBI:ARO2-His, which enhances CNGC14 abundance in the heterologous system^5^. As the source of AFB1, we used total protein extracts from *Arabidopsis thaliana* seedlings expressing pAFB1:ccvAFB1-mScarlet.

Fresh *N. benthamiana* leaves were ground using the CNGC14 extraction buffer (50 mM HEPESpH 7.5), 100 mM KCl, 100 mM MgCl₂, 100 mM sucrose, 5mM DTT and 0.5% Igepal), supplemented with EDTA-free protease inhibitor cocktail (Pierce, A32965) and 1 mM PMSF at 4 °C. Incubation was performed at 4 °C for 30 min at constant rotation, filtered in miracloth (475855, Millipore) followed by centrifugation at 10000 × g for 10 min at 4 °C to remove debris.

Arabidopsis seedlings were harvested, flash-frozen in liquid nitrogen, and homogenized in AFB1 extraction buffer (20mM HEPES pH6.8, 150 mM NaCl, 1mM EDTA, 1mM DTT, and 0.5% Tween 20) supplemented with 1 mM PMSF and cOmplete protease inhibitor cocktail (Roche). Incubation was carried out for 1 h at 4 °C, and clarified by centrifugation at 20,000 × g for 25 min at 4 °C.

For the pull-down reaction, total protein extracts from *N. benthamiana* leaves were first incubated with GFP-Trap Magnetic Particles M-270 (Chromotek), pre-equilibrated in CNGC14 extraction buffer, for 1.5 h at 4 °C with continuous rotation. After incubation, the magnetic particles were washed twice with CNGC14 extraction buffer and once with AFB1 extraction buffer. Subsequently, the beads were incubated with total protein extract from Arabidopsis seedlings in the presence of either 50 µM cvxIAA (treatment) or the corresponding volume of DMSO (mock) for 1 h at 4 °C with end-over-end rotation. Following the incubation, the beads were washed 4 times in the AFB1 extraction buffer supplemented with either 10 µM cvxIAA (treatment) or DMSO (mock).

Bound proteins were eluted by heating to 65 °C for 10 min in 2× Laemmli sample buffer (Bio-Rad). Samples were separated by SDS-PAGE and analyzed by western blotting using anti-GFP antibody (Roche 11814460001) for detection of CNGC14-GFP and anti-mCherry antibody (Agrisera AS16 3967) for detection of AFB1-mScarlet. The experiment was repeated twice with identical results in a fully biological and technical repeat.

#### DII pulldown experiment

DII-peptide pull-downs were performed as described previously^2^. In brief, seven-day-old seedlings of pTIR1:ccvAFB1-mScarlet, pAFB1:ccvAFB1-mScarlet and pTIR1:ccvAFB1-Y78F-mScarlet were harvested and homogenized for total protein extraction in the AFB1 extraction buffer. The N-terminally biotinylated CNGC14 C-terminal helix peptide (TSNVKPHFAATILASRFAKNTRRTAHKLK), CNGC14-FAFA peptide (TSNVKPHAAATILASRAAKNTRRTAHKLK), CNGC6 C-terminal helix peptide (AGGSPYSIRATFLASKFAANALRSVHKNR), and Aux/IAA7 DII peptide (AKAQVVGWPPVRNYRKN) were coupled to streptavidin–agarose (Millipore) beads following Kepinski et al.^9^. For pull-down reactions, conjugated beads were incubated with total protein extracts in the presence of either 50 µM cvxIAA or the corresponding volume of DMSO (mock) for 1 h at 4 °C with end-over-end rotation. The NaCl concentration was adjusted to 450mM for CNGC14, CNGC6 and CNGC14-FAFA, and 150 mM for Aux/IAA7 DII peptide in the pull-down reaction as well as in the following washing steps. Beads were washed three times in the AFB1 extraction buffer supplemented with 1 mM PMSF and cOmplete protease inhibitor cocktail (Roche) and either 10 µM cvxIAA (treatment) or DMSO (mock). Western blot for AFB1-mScarlet identification was performed using anti-mCherry antibody (Agrisera AS16 3967).

### Proximity labeling

The pAFB1:ccvAFB1-turboID-YFP or free pAFB1:Free-turboID-YFP lines were used. T3 generation lines with moderate expression levels were selected for the experiment. We followed the protocol by Arora^10^ for Arabidopsis cell cultures, which we optimized for Arabidopsis seedlings: Plants were grown on a vertical 1/2 MS medium covered with a sterile Uhelon 74T polyamide mesh with 85 μm pores (Silk & Progress) to facilitate root harvesting. Prior to the harvest, seedlings were treated for 1 hour at room temperature with 50 μM Biotin (Sigma-Aldrich) plus either 500 nM cvxIAA (treatment) or DMSO (mock) in water. Roots were cut, weighed, and homogenized in liquid nitrogen. Samples were incubated for 1h on a rotating device in the extraction buffer EB (100 mM Tris-HCl pH 7.5, 2% (w/v) SDS, 8 M urea), centrifuged 2 x 20 min at a maximum speed, and the supernatant was filtered through a RC25 0.45 μm Syringe filter. Excess biotin was removed on PD10 desalting column pre-equilibrated with the Binding buffer BB (100 mM Tris-HCl pH 7.5, 2% (w/v) SDS, 7.5M urea). Proteins were eluted by the addition of 3.5 ml of EB and collected into the 5 ml LoBind tubes containing equilibrated Streptavidin-Sepharose beads (100 μl/mg of fw sample, Cytiva), and incubated overnight on a rotating device. The next day, Streptavidin-Sepharose beads were gently spun at 450 g for 1 min and transferred (as a 500 μl bead slurry) onto a 0.35-μm column bottom filter. 5 washing steps with 800 μl of BB were followed by incubation in High salt buffer (1M NaCl, 100 mM Tris-HCl pH 7.5) for 30 min. After 2 washing steps with miliQ water and 4 washing steps with 50 mM Tris pH 8.0, excess liquid was removed, and wet beads were frozen in liquid nitrogen prior to MS-MS analysis.

### Proteomics

#### Protein Digestion

Beads from immunoprecipitation were mixed with 1 M TEAB (Triethylammonium bicarbonate) in 20% SDC (sodium deoxycholate), 100 mM TCEP (Tris(2-carboxyethyl)phosphine), and 500 mM CAA (chloroacetamide) and shaken for 30 min at 60°C. After incubation, IP beads were digested in 50 mM TEAB with 0.5 µg of trypsin-lysC protease mix (Thermo Fisher Scientific) at 37 °C overnight. After digestion, samples were centrifuged, and the supernatant was acidified with TFA to a final concentration of 1%. The supernatant was washed 3times with ethyl acetate, and the residual ethyl acetate was evaporated. Samples were acidified with TFA to a final concentration of 1% and the peptides were desalted using in-house-made stage tips packed with C18 disks (AttractSPE, Affinisep, France) according to Rappsilber et al.^11^

#### nLC-MS 2 Analysis

Nano Reversed phase column (Ion Opticks, Aurora Ultimate TS 25×75 C18 UHPLC column) was used for LC/MS analysis. Mobile phase buffer A was composed of water and 0.1% formic acid. Mobile phase B was composed of acetonitrile and 0.1% formic acid. Samples were loaded onto the trap column (C18 PepMap100, 5 μm particle size, 300 μm x 5 mm, Thermo Scientific) for 4 minutes at a flow rate of 18 μl/min. The loading buffer was composed of water, 2% acetonitrile, and 0.1% trifluoroacetic acid. Peptides were eluted with Mobile phase B gradient from 4% to 35% B in 60 min.

Eluting peptide cations were converted to gas-phase ions by electrospray ionization and analyzed on a Thermo Orbitrap Ascend (Thermo Scientific) by data-dependent approach. Survey scans of peptide precursors from 350 to 1400 m/z were performed in Orbitrap at 120K resolution (at 200 m/z) with a 100 % ion count target. Tandem MS was performed by isolation at 1,6 Da with the quadrupole, followed by HCD fragmentation with normalized collision energy of 30 % and 10 ms activation time.

Fragmentation spectra were acquired in ion trap with scan rate set to Rapid. The MS2 ion count target was set to 150 % and the max injection time was 75 ms. Only those precursors with charge state 2–6 were sampled for MS2. The dynamic exclusion duration was set to 30 s with a 10ppm tolerance around the selected precursor and its isotopes. Monoisotopic precursor selection was turned on. Cycle time was set to 1.5 s.

#### Proteomic sample preparation

Proteins bound to beads were eluted by incubation for 10 min at 65 °C with continuous shaking in 80 µL of 2% sodium deoxycholate (SDC) in 100 mM triethylammonium bicarbonate (TEAB). Proteins were reduced with 10 mM TCEP (2-carboxyethyl phosphine) and alkylated with 25 mM chloroacetamide at 95 °C for 5 min.

Proteolytic digestion was performed by adding 0.5 µg of Lys-C/trypsin, followed by incubation for 2 h, after which an additional 0.5 µg of Lys-C/trypsin was added. Samples were then incubated overnight at 37 °C.

Digestion was stopped by acidification with trifluoroacetic acid (TFA) to a final concentration of 1%. SDC was removed by liquid–liquid extraction as described previously^12^, and samples were evaporated to dryness. The resulting peptides were reconstituted in 15 µL of loading buffer and analyzed by LC-MS/MS.

Liquid chromatography separation was performed using a Vanquish Neo UHPLC system (Thermo Fisher Scientific) operated in trap-and-elute mode. Samples were loaded onto a trap column (C18 PepMap100, 5 µm particle size, 300 µm × 5 mm, Thermo Scientific) under combined flow/pressure control (maximum flow 100 µL/min or maximum pressure 800 bar). The loading buffer consisted of water with 2% acetonitrile and 0.1% formic acid. Mobile phase A consisted of water with 0.1% formic acid, and mobile phase B consisted of acetonitrile with 0.1% formic acid. Peptides were eluted using a linear gradient from 2% to 35% mobile phase B over 32 min.

Peptides were separated on a nano reversed-phase column (Aurora Ultimate TS, 25 cm × 75 µm ID, 1.7 µm particle size, IonOpticks) operated at a flow rate of 0.4 µL/min and analyzed online by an Orbitrap Astral Zoom mass spectrometer (Thermo Fisher Scientific) using a data-independent acquisition (DIA) approach. Eluting peptide cations were ionized by electrospray ionization in positive mode with a spray voltage of 1.6 kV and an ion transfer tube temperature of 280 °C.

MS1 scans were acquired in the Orbitrap over an m/z range of 380–980 at a resolution of 120,000 with the following settings: RF lens 40%, maximum injection time 3 ms, and normalized AGC target 500%. DIA MS2 scans were acquired in the Astral analyzer using 2 Da isolation windows, with a normalized AGC target of 500%, maximum injection time of 3 ms, and pre-accumulation enabled. Fragment ion spectra were acquired over an m/z range of 150–2000 using higher-energy collisional dissociation (HCD) with a normalized collision energy of 25%^13^.

#### MS data processing

The acquired data were analyzed using Spectronaut (version 20.2.250922.92449)^13^. Data were processed using the directDIA+ (Deep) workflow with normalization enabled. Searches were performed against an *Arabidopsis thaliana* database (one protein per gene, 27448 entries) downloaded on 20.10.2025 and supplemented with common contaminants and sequence of bait protein.

Search parameters included trypsin enzyme specificity, carbamidomethylation of cysteine as a fixed modification, and protein N-terminal acetylation and methionine oxidation as variable modifications. Peptide-and protein-level false discovery rates (FDRs) were controlled at 1%. Further filtering of data and statistical analysis was performed using Perseus software^14^.

#### Data analysis

All data were analyzed and quantified with the MaxQuant software (version 2.4.13.0)^15^. The false discovery rate (FDR) was set to 1% for both proteins and peptides and we specified a minimum peptide length of seven amino acids. The Andromeda search engine was used for the MS/MS spectra search with the *Arabidopsis thaliana* database (downloaded from Uniprot in March 2024, containing 27 449 entries). Enzyme specificity was set as C-terminal to Arg and Lys, also allowing cleavage at proline bonds and a maximum of two missed cleavages. Carbamidomethylation of cysteine was selected as fixed modification and N- terminal protein acetylation and methionine oxidation as variable modifications. Data analysis was performed using Perseus 1.6.15.0 software^14^. To create the volcano plot in Fig. 1F despite the NaN values in the negative control and mock treatment, the missing values were imputated by values corresponding to the detection limit of the system.

### High-resolution imaging

Imaging was performed using a vertical-stage Zeiss Axio Observer 7 microscope^16^ equipped with a Yokogawa CSU-W1-T2 spinning disk unit (50 μm pinholes) and a VSHOM1000 excitation light homogenizer (Visitron Systems). Images were acquired using VisiView software (Visitron Systems, v.4.4.0.14). For imaging, seedlings were mounted in microscopy chambers and gently covered with solidified growth medium supplemented with 100 nM IAA or 500 nM cvxIAA for transgenic lines expressing nAFB1 or ccvAFB1, respectively. Mock treatments used 0.5 μL/mL DMSO or 0.1 μL/mL ethanol. Imaging was performed within 30 minutes of treatment using a Zeiss Plan-Apochromat ×100/1.46 oil immersion objective with 488 nm excitation, 500-550 nm emission for GFP; 515 nm excitation, 520-570 nm emission for mVenus; 561 nm excitation, 582-636 nm emission for mScarlet. Fluorescence signals were captured using a PRIME-95B back-illuminated sCMOS camera (1,200 × 1,200 pixels; Photometrics). For colocalization, GFP and mScarlet channels were imaged sequentially. Acquired images of the epidermal surface in the root elongation zone were processed using FIJI^7^, and automatic identification of nAFB1 dots was performed using the TrackMate plugin v7.9.2^17^.

### FRET-FLIM imaging

For FRET-FLIM, either roots of 5-day-old *A. thaliana* seedlings or leaves of *N. benthamiana* expressing the proteins of interest were imaged. Immediately before imaging, the seedlings were transferred to 1/2 MS medium containing treatment (100 nM IAA or 500 nM cvxIAA) or mock (EtOH or DMSO, respectively) and then as described previously. Similarly, the leaves of *N. benthamiana* were infiltrated with MiliQ H2O containing either treatment or mock immediately before imaging.

The microscope used was a confocal laser scanning microscope Zeiss LSM880, with an apochromatic 40x/1.2 objective, running ZEN Blue acquisition software. mEGFP was excited with a pulsed tunable multi-photon laser Chameleon Ultra II (Coherent) set to a wavelength of 930 nm and a laser intensity of 4%, and a 525/50 filter set for eGFP emission. An area containing cells with good expression of the protein(s) of interest was selected using this setup. Lifetime data was acquired with a Becker-Hickl card SPC-150, a hybrid detector for Time-Correlated Single-Photon Counting, and the SPCM64 data acquisition software running on a separate computer.

Images were recorded at 512 x 512 pixels, with a time resolution of 1024 time channels per pixel and an acquisition time of approximately 4 minutes per image. The data were fitted to a single exponential parameter with an incomplete multi-exponential decay model using the SPCImage v8.5 software while considering the goodness of the fit (χ2 ∼ 1). An incomplete decay model was used because some residual fluorescent signal from the previous laser pulse was still present at the start of the next pulse. The fitting was performed using the Instrument Response Function (IRF), which was recorded using the MeOH-urea sample.

For both samples from roots of *A. thaliana* and leaves of *N. benthamiana*, the images were processed by selecting a population reasonably close to the universal semicircle on the phasor plot to exclude signals heavily contaminated with autofluorescence (e.g., chloroplasts in leaves). For the root samples from *A. thaliana*, mean fluorescent lifetimes and colored lifetime images were calculated in SPCImage. For the samples from leaves of *N. benthamiana*, regions of interest where neither of the fluorophores was present in excess were selected, and mean lifetimes were extracted from those regions only.

### Real-Time Imaging of Ca^2+^ Fluxes in Roots

Ca^2+^ imaging was performed using a microfluidic chip system^18^, mounted on the vertical imaging system (described above). Five-day-old Arabidopsis seedlings were gently transferred into the microfluidic chip. Roots were first perfused with a mock solution composed of ½ MS medium, 2 µg/mL dextran labeled by tetramethylrhodamine (D1816, Invitrogen), 0.5 µL/mL DMSO, adjusted to pH 5.8, at a constant flow rate of 3 µL/min for 15–30 minutes to establish baseline conditions.

Subsequently, the roots were treated with a treatment solution containing ½ MS medium, 1 µg/mL TMR-dextran, and 500 nM cvxIAA, also at pH 5.8 and the same 3 µL/min flow rate. Imaging was conducted over a total period of 15 minutes, consisting of 5 minutes of mock perfusion followed by 10 minutes of treatment, with a 7 s acquisition interval. Throughout the imaging session, the pressure within the microfluidic chip was maintained below 90 mbar to avoid mechanical stress on the roots. Acquired images were analyzed in FIJI^7^ to measure GCaMP3 fluorescence in epidermal cells of the root transition zone.

### Transcriptomic analysis

For RNA seq, 5-day-old seedlings of pTIR1:ccvTIR1-mScarlet, pAFB1:ccvAFB1-mScarlet, and Col-0 Arabidopsis lines were treated with 500 nM cvxIAA^19^ or mock (DMSO) in 1/2MS with 1% sucrose for 1 hour; RNA was isolated from roots using Plant Total RNA Mini Kit, (Favorgen). The quality of isolated RNAs was established by agarose electrophoresis and determination of RIN using Bioanalyzer (Agilent). Strand-specific cDNA libraries were constructed from polyA-enriched RNA and sequenced on the Illumina NovaSeq 6000 platform. The sequencing resulted in at least 20 million 150 bps long read pairs. Rough reads were quality-filtered using Rcorrector^20^ and Trim Galore (https://www.bioinformatics.babraham.ac.uk/projects/trim_galore/) scripts using default parameters. Levels of transcript abundances were determined using Salmon^21^ with parameters --posBias, --seqBias, --gcBias, --numBootstraps 30. Reference index was built from TAIR10 cds library, version 20101214, with added coding sequences of ccvAFB75-mScarlet and ccvTIR1-mScarlet. Statistical evaluation and quality control of data analysis was done using sleuth R package, version 0.30.2^22^.

### Structural modeling

A local version of AlphaFold3^23^, provided by e-INFRA CZ, which enables the use of custom ligands, was employed for modelling with mostly full-length protein sequences as input. Five different seeds were used for each complex. The best-scoring models were selected based on the predicted alignment error (PAE) plots. UCSF ChimeraX^24^ was used to visualise the protein complexes, PAE plots, and create figures.

## Acknowledgements

LCMS analyses were performed in the Laboratory of Mass Spectrometry at Biocev research center; Faculty of Science, Charles University. The authors thank Ramilla Brikova for help and support and George Caldarescu for the help with the proximity labeling experiments.

## Diversity, equity & inclusion statement

Members of our author team bring diverse identities, including representation from LGBTQIA+ communities.

## Data and materials availability

All the data are included either in the main manuscript or in supplementary material. The materials will be available upon request.

## Additional information

**Table S1.** (separate file) Transcriptome of ccvTIR1 and ccvAFB1 in response to cvxIAA

**Table S2.** (separate file) List of the top enriched proteins identified by AFB1 pulldown from Arabidopsis roots and whole seedlings.

**Table S3.** (separate file) List of the proteins identified in the ccvAFB1 proxiome.

**Table S4.** (separate file) List of generated constructs and transgenic lines.

**Table S5.** (separate file) List of primers used in the study.

**Table S6.** (separate file) List of the proteins co-immunoprecipitated with CNGC14-GFP.

**Movie S1. ccvAFB1 receptor triggers a cytosolic Ca^2+^ transient in response to cvxIAA ligand.**

Simultaneous imaging of roots expressing the GCaMP3 calcium reporter in Col-0 (WT, left root) or ccvAFB1-mScarlet (magenta, right root) grown in a microfluidic device. The cvxIAA ligand (500 nM) triggers a visible increase in GCaMP3 fluorescence in the ccvAFB1-mScarlet expressing root but not in WT. The cvxIAA application is visualized by the decrease in the medium fluorescence tracer intensity and indicated in the movie at time point 0s. Scale bar: 50 μm, time interval: 7s.

**Movie S2. ccvAFB1mVenus forms foci on plasma membrane in response to the cvxIAA ligand.**

Time series of the surface of ccvAFB1-mVenus expressing root epidermal cell imaged with a spinning disk microscope shows the presence of the receptor in foci that dwell on the plasma membrane upon 500 nM cvxIAA treatment. Scale bar: 5 μm, time interval: 1.5s.

**Movie S3. ccvAFB1 mScarlet colocalization with CNGC14-GFP.**

Time-series imaging of root epidermal cells expressing ccvAFB1-mScarlet (left, magenta) and CNGC14-GFP (center, green), with merged channels (right) shown on the right. Both channels were imaged sequentially. Scale bar: 5 μm; time interval: 2.5 s.

**Movie S4. ccvAFB1 mScarlet colocalization with ARO2-GFP.**

Time-series imaging of root epidermal cells expressing ccvAFB1-mScarlet (left, magenta) and ARO2-GFP (center, green), with merged channels shown on the right. Both channels were imaged sequentially. Scale bar: 5 μm; time interval: 2.5 s.

**Movie S5. The ccvAFB1 with mutation in putative AC and GC domains is able to induce Ca^2+^ transient in response to the ligand.**

Simultaneous imaging of roots expressing the GCaMP3 calcium reporter in Col-0 (WT, left root) or ccvAFB1dAC/GC-mScarlet (magenta, right root) grown in a microfluidic device. The cvxIAA ligand (500 nM) triggers a visible increase in GCaMP3 fluorescence in the ccvAFB1dAC/GC-mScarlet expressing root but not in WT. The cvxIAA application is visualized by the decrease in the medium fluorescence tracer intensity and indicated in the movie at time point 0s. Scale bar: 50 μm, time interval: 7s

## Supplementary Figures

**Fig. S1.**
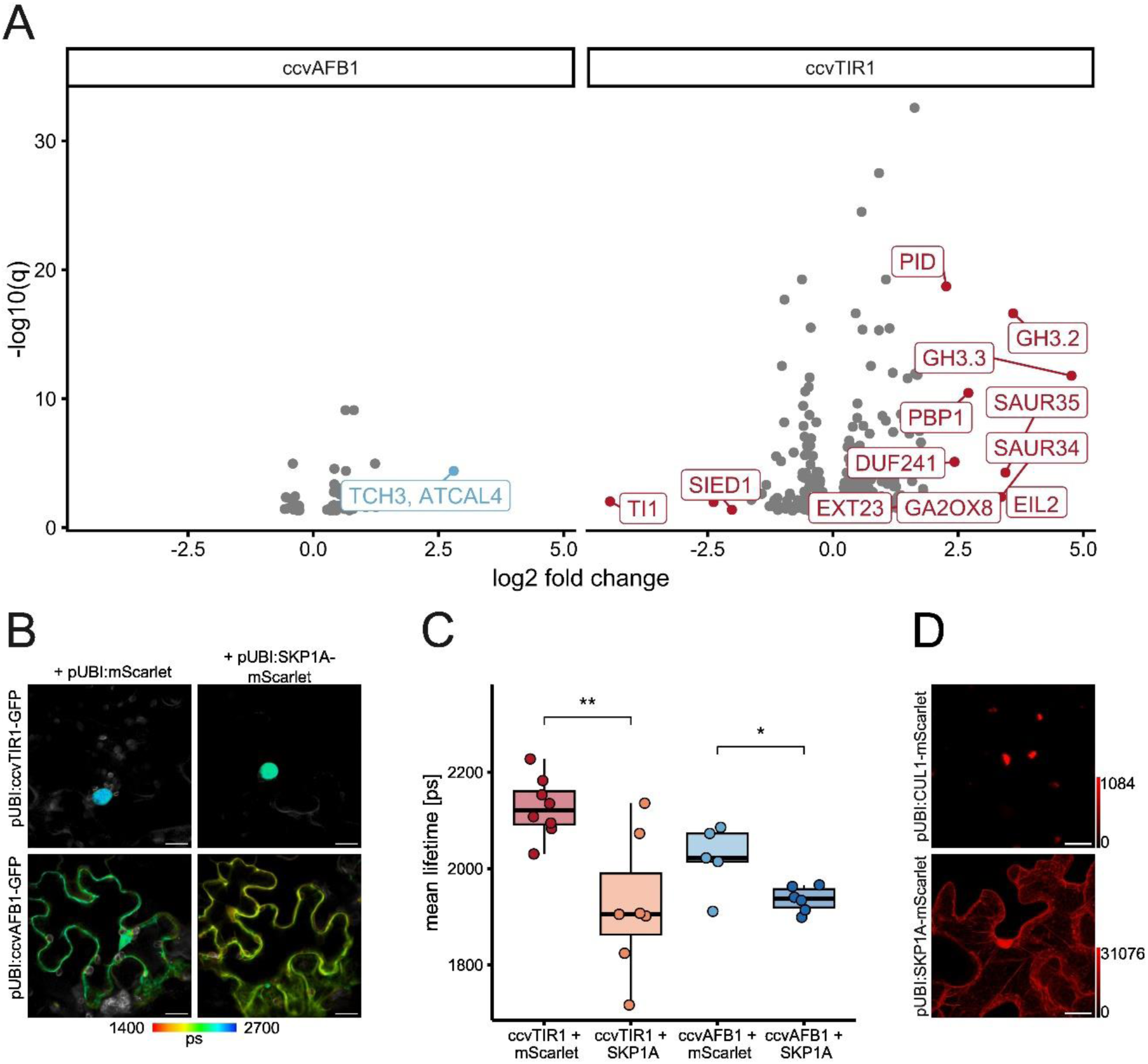
A) ccvAFB1 does not trigger auxin-induced gene expression. Volcano plot showing differentially expressed genes in the ccvTIR1 (red) and ccvAFB1 (blue) lines treated with 500 nM cvxIAA or mock (DMSO). Each point represents a gene, plotted according to the log2 fold change (x-axis) and-log10 adjusted p value (y-axis). Genes significantly upregulated after cvxIAA treatment are shown on the right, while significantly downregulated genes are shown on the left. Non-significantly changed genes are clustered near the center. Statistically significant DEGs at log2FC>2 are highlighted and annotated. B) fluorescence lifetime images of tobacco leaf epidermal cells and C) lifetime quantification of ccvAFB1-GFP and ccvTIR1-GFP with SKP1A-mScarlet and free mScarlet as a negative control. The pseudocolor represents the mean fluorescence lifetime, the intensity represents the photon count. The pseudocolor key shows the scale of the fluorescent lifetimes. Statistical comparisons in C) are done pairwise, comparing the effect of co-expressing the receptor (ccvTIR1-GFP or ccvAFB1-GFP) with either free mScarlet (negative control) or SKP1-mScarlet. Scale bars: 20 μm. D) CUL1-mScarlet localizes exclusively to nuclei, while SKP1A-mScarlet is present in nuclei and cytoplasm in tobacco leaf epidermal cells. Scale bars: 20 μm.

**Fig. S2.**
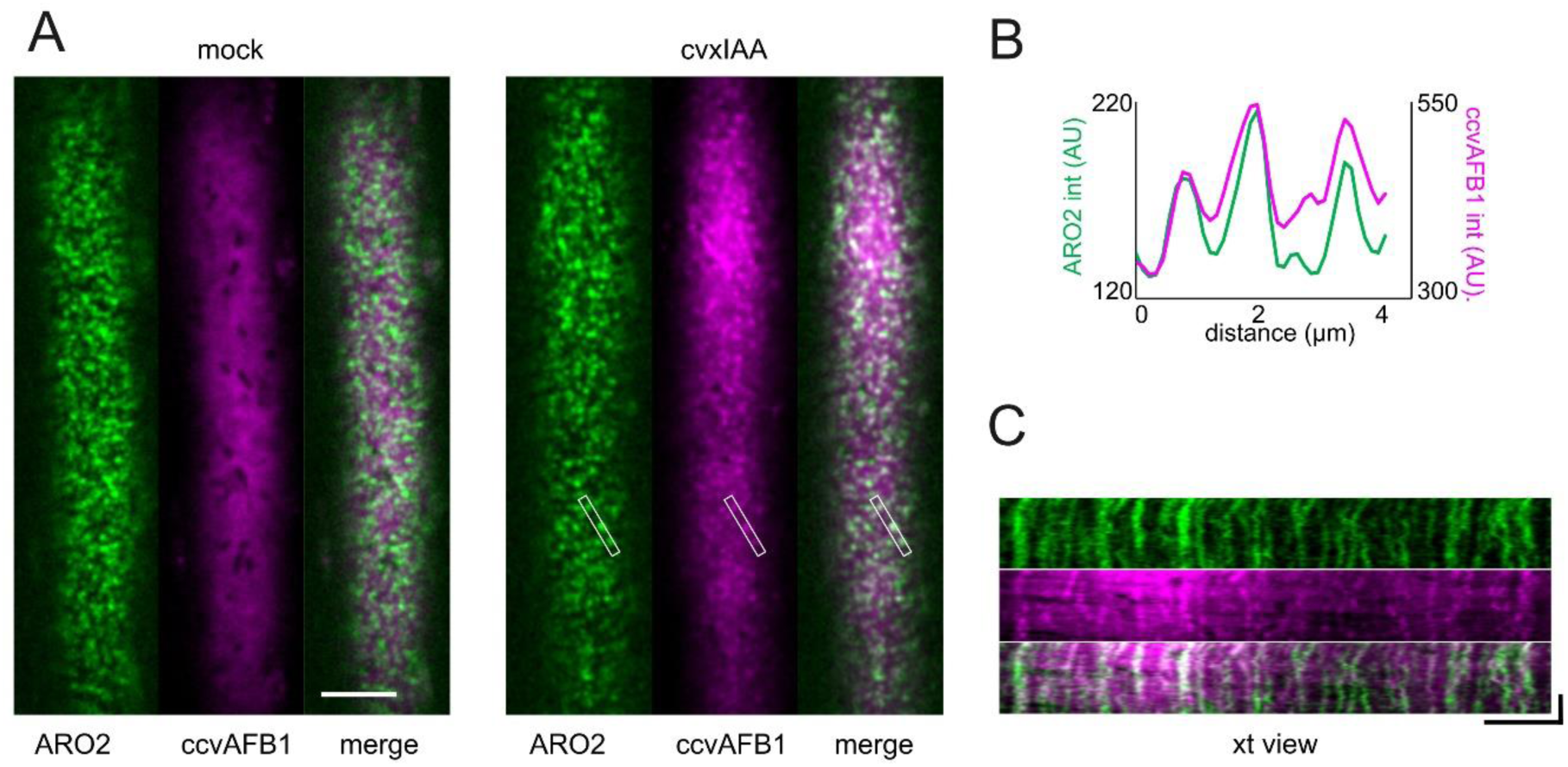
A-C) cvxIAA (500 nM) induced the colocalization of ccvAFB1-mScarlet with ARO2-GFP in root epidermal cells. A) single cell surfaces of Arabidopsis roots expressing ARO2-GFP (green) and ccvAFB1-mScarlet (magenta). B) A plot of CNGC14-GFP and ccvAFB1 fluorescence intensities along the line highlighted in (A). C) The kymograph (time-space view, xt view) shows the colocalization of ccvAFB1-mScarlet (magenta) with ARO2-GFP (green) in PM foci upon cvxIAA treatment. Horizontal dimension represents the x-dimension, while horizontal dimension represents t-time. Scale bars in all figures = 5 μm; vertical scale bar in(C) = 60 s.

**Fig. S3.**
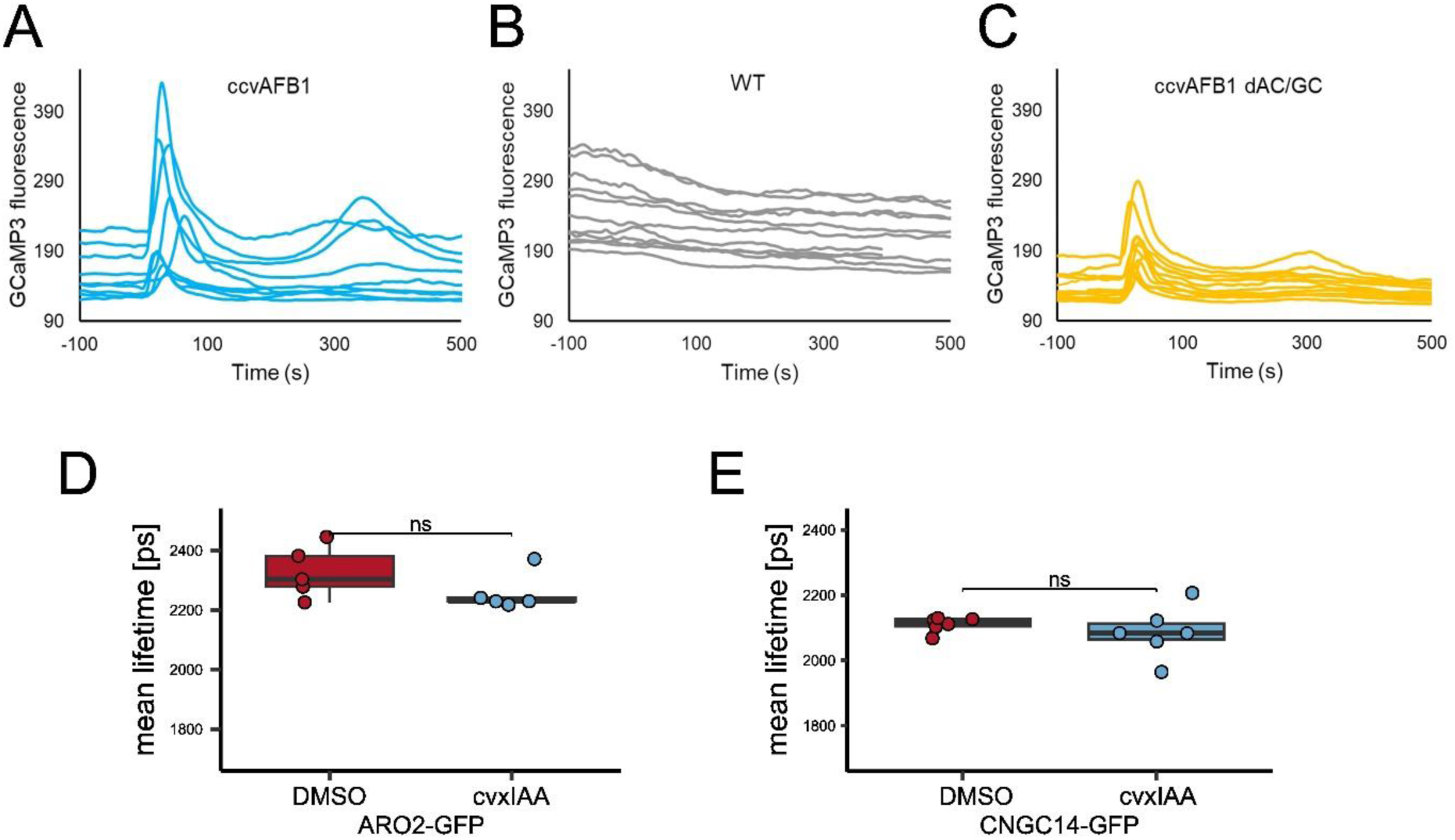
A-C) Raw non-normalized intensities of GCaMP3 fluorescence in response to 500 nM cvxIAA treatment in ccvAFB1-mScarlet (A), WT (B), and ccvAFB1dAC/GC-mScarlet(C) lines. Intensity was measured in epidermal cells of the root elongation zone. D) Quantification of FLIM images of ARO2-GFP and E) CNGC14-GFP in Arabidopsis root epidermal cells. Each dot represents the mean lifetime of a single root imaged under the respective conditions.

**Fig. S4.**
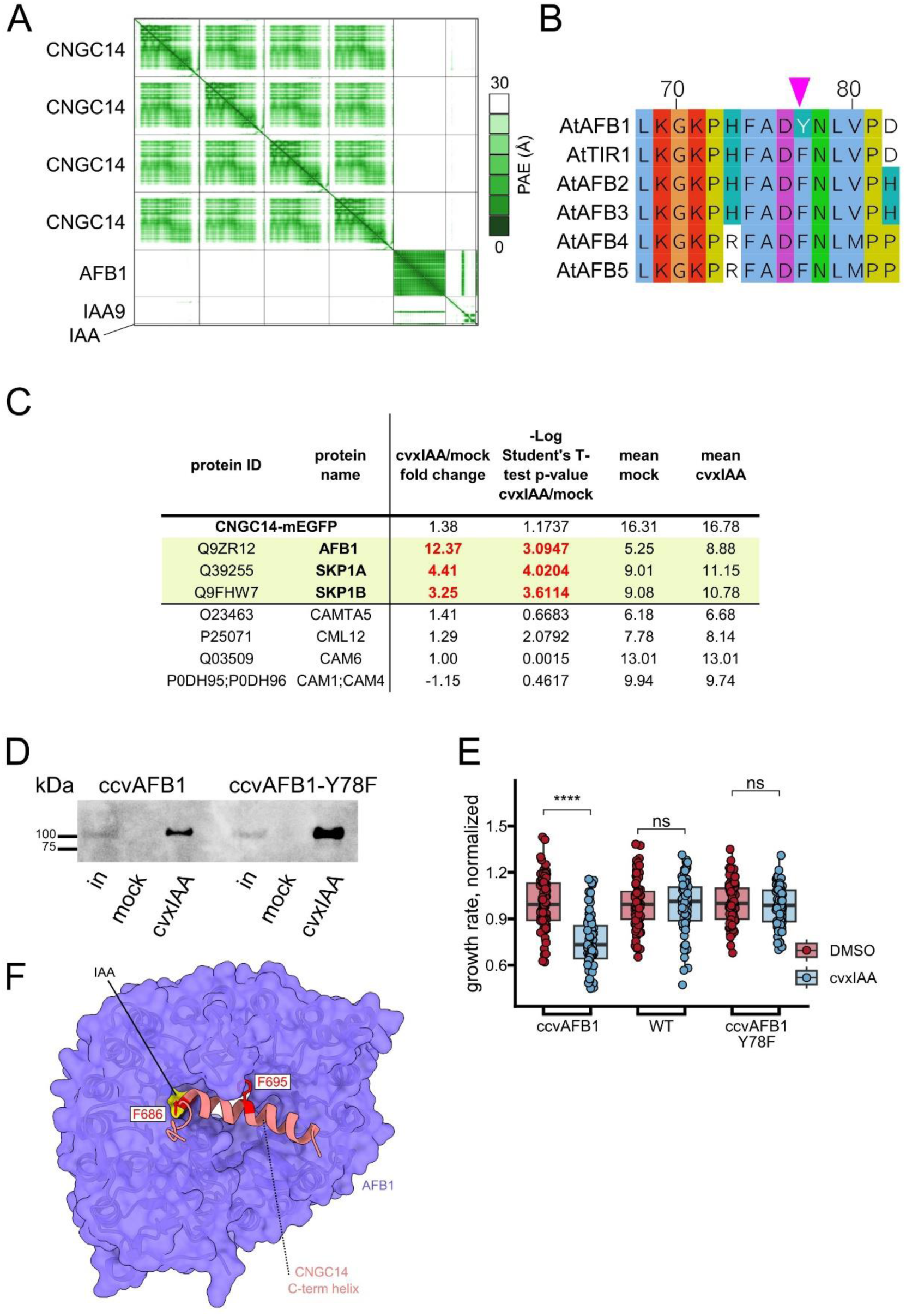
A) The predicted alignment error (PAE) plot for the complex composed of IAA, IAA9, AFB1, and the CNGC14 tetramer. AlphaFold3 predicts the IAA-mediated interaction between AFB1 and IAA9 with high confidence. B) Unlike the other Arabidopsis TIR1/AFB receptors, AFB1 possesses a single amino acid substitution (F78Y) in its auxin binding pocket. C) Mass-spectrometry analysis of the proteins co-immunoprecipitated with CNGC14-GFP reveals only 3 targets significantly enriched after cvxIAA treatment (significant fold change highlighted in red numbers). Mean values for both mock and cvxIAA treatment are calculated from three technical replicas. The bottom part of the table shows all calmodulins present in the dataset. D) In vitro pull-down assays of ccvAFB1-mScarlet and ccvAFB1-Y78F-mScarlet by IAA7-DII peptide co-incubated with either mock (DMSO) or 50 μM cvxIAA. Input is shown (in); ccvAFB1-mScarlet detected by anti-mCherry antibody. E) ccvAFB1-Y78F fails to inhibit root growth in response to auxin. Normalized growth rates of pAFB1:ccvAFB1, Col-0 (WT), and p35S:ccvAFB1-Y78F seedlings transferred on either control DMSO or 500 nM cvxIAA treatment for four hours. F) The top view of the interaction interface between C-terminal helix of CNGC14 and AFB1 highlights the F686 and F695 residues that are in contact with auxin and the auxin-binding pocket, respectively. An AlphaFold3 prediction of the IAA-dependent interaction between AFB1 (blue) and the C-terminal helix of CNGC14 (amino acids 683-708, light red).

